# Navigating the pangenome coordinate system with Shredtools

**DOI:** 10.64898/2026.07.03.736354

**Authors:** Vikram S. Shivakumar, Ben Langmead, Human Pangenome Reference Consortium

**Affiliations:** Department of Computer Science, Johns Hopkins University

## Abstract

Existing notions of pangenome coordinates rely on hard-to-compute multiple sequence alignments. On the other hand, pangenome-wide exact unique matches (multi-MUMs) can be computed efficiently, and represent conserved stretches of columns in the underlying MSA. We introduce Shredtools, which uses multi-MUMs as pangenome waypoints and allows for sophisticated queries in pangenome coordinates. Its primary query is `extract`, which takes an interval of one sequence and extracts the smallest window containing it that is syntenic pangenome-wide. Shredtools’ `extract` query can extract a gene region from 476 human genomes in half a second. Other queries help to refine these results, by finding local exact matches to improve the density of multi-MUM coverage (“enhance”) and by selectively discarding sequences to improve the precision of the syntenic region (“zoom”). The Shredtools web interface (available at https://vikshiv.github.io/shredtools) allows for client-side handling of `extract` queries with index queries handled via simple and fast HTTP Range requests, simplifying usage and enabling pangenome-scale discoveries.

## Introduction

Pangenome collections are growing rapidly, accumulating new high-quality genomes from across the tree of life. We use the term *genome* throughout to refer to each individual member of a pangenome collection. When using a single-genome reference, existing tools can straightforwardly extract genomic intervals corresponding to, e.g., annotated genes. For pangenomes, this is less straightforward, and the relevant methods and tools are still being developed.

To date, several approaches have been proposed for defining pangenome representations and associated coordinate systems. The most widely adopted approaches represent the pangenome as a graph, where vertices and edges encode relationships among homologous genomic segments [1], [2], [3]. The graph structure is derived from a multiple sequence alignment (MSA), or an approximation thereof. Graph-based representations permit genomic loci to be described relative to the graph itself or relative to one of the genomes threaded through the graph. Features that are mapped onto the graph can also be lifted (surjected) to other genomes in the graph.

While this system is workable overall, the whole-genome MSAs underlying these graphs are extremely hard to compute. Computing an MSA is already infeasible for today’s pangenomes, such as the Human Pangenome Reference Consortium release 2 [4]. Even for pangenomes where MSAs have been computed, such as the HPRC release 1 [5], building it required masking some of the most repetitive sequences [5], which are also the sequences that long-read sequencing and T2T workflows were needed to resolve.

Alternatively, pangenome coordinates can be built by composing many pairwise whole-genome alignments. In such a system, users can analyze data relative to one linear reference genome, then subsequently transfer, or “lift,” to another linear genome when necessary. impg [6], for example, projects intervals across genomes from whole-genome pairwise alignments and underlies the PGGB pipeline [7]; similar liftover ideas appear in alignment-based frameworks such as levioSAM2 [8]. While these methods can work for moderate-sized pangenomes, the need to compute alignments over all genome pairs becomes intractable for large pangenomes. Computing pairwise alignments over only some of the pairs may require lifting across a sequence of intermediate linear coordinate systems before arriving at the destination, leading to reduced fidelity for genome pairs separated by more intermediate genomes.

This work takes a different approach, building on the notion of pangenome way-points or markers: sequences that are shared across the collection but unique within each genome. This approach has precedents; ntsynt, for instance, uses shared minimizers to split genomes and detect synteny within a pangenome [9]. A closer analogue is AGC [10]. AGC uses unique *k*-mers, i.e. substrings of fixed length *k* that are shared across the collection yet unique in each genome, to divide the pangenome into components that are amenable to compression. Computing markers is far more efficient than computing an MSA, and, as we argue here, markers can suffice for pangenome queries. Prior marker-based methods, however, rely on one or more fixed length parameters: the *k*-mer length *k*, and minimizer window size. Those must be chosen in advance, even though the best choice varies with the size and diversity of the collection.

We use multiway maximal unique matches (multi-MUMs) as markers. These are maximal substrings that occur exactly once in every genome in the collection. We recently introduced Mumemto [11], which computes multi-MUMs efficiently even for the largest pangenomes, and showed that they mark conserved, collinear columns of the underlying MSA. Crucially, multi-MUMs require no length parameter: because they are maximal and variable-length, a single multi-MUM index simultaneously captures markers of every length. Multi-MUMs thus generalize fixed-length waypoints such as unique *k*-mers and minimizers, and we show below that they cover more conserved sequence than the best attainable *k*-mer marker set at any single *k*, giving a faithful and parameter-free picture of the pangenome coordinate system. Multi-MUMs have also proven useful as building blocks: we previously used them to construct a linearized pangenome graph [11], and, in related work, to mark collinearity within an *r*-index, improving read classification against pangenome references without a full MSA [12].

Multi-MUMs are also readily mergeable: MUMs computed over partitions of a pangenome can be combined into the multi-MUMs of their union without repeating the most expensive steps. This brings dynamism to the coordinate system; newly released genomes can be folded in rapidly.

Here, we introduce Shredtools, a toolkit for navigating and manipulating pangenomes through their multi-MUM coordinate system. Given Mumemto-computed multi-MUMs, Shredtools can extract the sequences syntenic to a region of interest, divide a pangenome into collinear local homologous regions for downstream computation, and refine the co-ordinate system toward a full multiple alignment by finding additional local matches. Because multi-MUMs are efficient to both compute and update, Shredtools scales to the largest available pangenomes, letting users retrieve local alignments and explore pangenome variation without first constructing an MSA or graph.

## Results

### Pangenomes used

For human pangenome applications, we used releases 1 and 2 from the Human Pangenome Reference Consortium (HPRC) [4, 5], which we call HPRCr1 (N=92) and r2 (N=476) respectively. We also used the following panel of T2T assemblies: CHM13 [13], YAO [14], HG002 [15], I002C [16], CN1 [17], KOREF1 [18], KSA001 [19], and PAN027 [20]. Note that these human pangenome collections consist of many diploid assemblies, and that we are using the term “genome” to refer to individual haplotypes from the collection.

### Multi-MUMs partition pangenomes better than k-mers

The Shredtools `shred` function splits a pangenome into collinear homologous regions using multi-MUMs as dividers, enabling downstream tasks such as graph construction to be parallelized over independent local subsets. Marker coverage determines the granularity of this partition: any stretch of genome not spanned by a marker becomes a single unsplit-table shred, so higher coverage yields finer-grained, more useful subsets. While *k*-mers can also serve as dividers [10], their coverage depends on the choice of *k*. We quantified this by computing, for each *k*, the fraction of CHM13 covered by unique *k*-mer markers across subsets of increasing size drawn from HPRCr2, comparing to multi-MUM coverage. Figure 1 shows that unique *k*-mer coverage is strictly below multi-MUM coverage for every collection size and every choice of *k*. Furthermore, the coverage-maximizing *k* decreases as the pangenome grows (from *k* = 67 at *N* = 10 to *k* = 37 at *N* = 400), confirming that no fixed length is optimal across collection sizes and that variable-length markers are necessary for consistent partitioning resolution.

**Figure 1:**
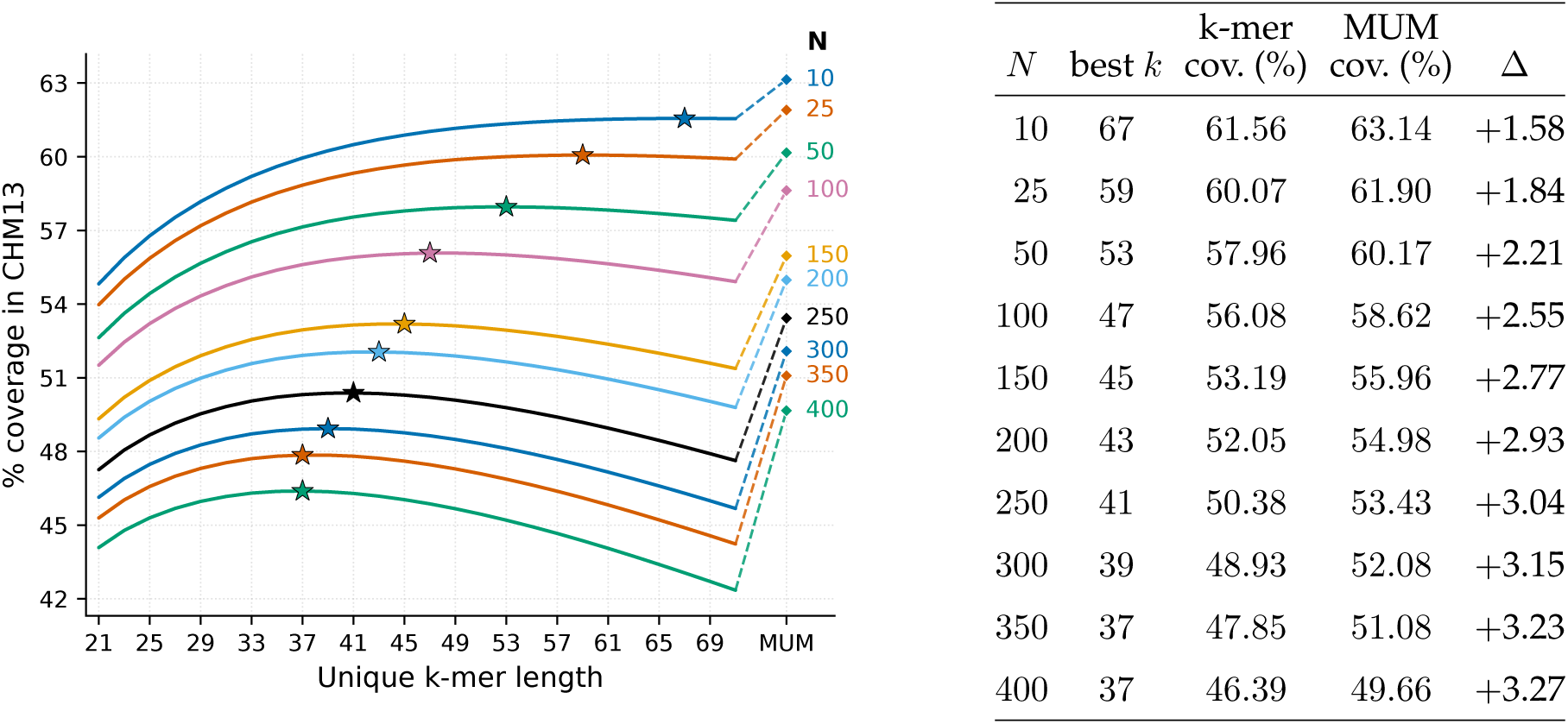
The choice of “best” *k*, which maximizes coverage of unique *k*-mers, varies with pangenome size. Multi-MUM coverage, using variable-length markers, yields optimal coverage of unique, collinear exact matches across all pangenome sizes, motivating their use as a common coordinate system.

### Shredtools extract is an efficient pangenome-coordinate query

In single-reference genomics, a basic and common operation is the interval query: retrieve all features (genes, etc.) overlapping a given locus. BEDTools [21] and `samtools view` make these queries routine, and many downstream analyses begin there. Pangenomics lacks an equivalent primitive: asking what sequence is present at a locus of interest *across all genomes in a collection* currently requires either a precomputed MSA, a pangenome graph, or a chain of pairwise liftover steps. Shredtools’ `extract` command fills this gap. It is roughly the pangenome analogue of `bedtools getfasta`: given any interval in any one genome, it retrieves the syntenic region from every other genome in the collection, using collinear multi-MUMs as coordinate anchors. Because multi-MUM indices are fast to build and compact to store, this query scales to the largest available pangenomes. To accelerate `extract`, we developed the `bumbl.bi` index, a data structure that enables rapid, memory-efficient retrieval of collinear multi-MUMs in a local region. `bumbl.bi` is described further in Methods.

We benchmarked `extract` against impg [6], which supports a similar query. The impg tool uses pairwise alignments and implicit interval trees to project aligned coordinates across the pangenome. For comparison, we generated an impg index and multi-MUMs across 92 chr19 assemblies from HPRCr1. impg involves all-pairs whole-genome alignments; as such, we were unable to compute a comparable index over whole assemblies in HPRCr1 or the larger HPRCr2 collection (N=476), both of which Shredtools was able to index and query.

Figure 2 compares runtime and peak memory for constructing Shredtools and impg indexes over N=92 chr19 genomes with different multithreading configurations. The Shredtools index was computed using multi-MUMs from Mumemto, while the impg index was built over alignments from wfmash [22]. Shredtools index construction was ∼ 86× faster than impg at 48 threads. Run serially in a single thread to minimize memory usage, Shredtools was 3.8× faster and used 20× less memory. In practice, impg operates over a sparse sampling of the pairwise comparison space. We ran wfmash on just 6% of pairwise alignments using 48 threads (“sparser impg”; figure 2). Compared to this configuration (using 48 threads), Shredtools (using 1 thread) was still 2× faster while using 15× less memory. The Shredtools index was a binary `bumbl` file containing multi-MUMs and the associated `bumbl.bi` index. This was 252.4 MB on disk. The corresponding impg index (including all PAF alignments + index) took 10.1 GB (40× larger), and 630 MB for the sparser configuration (2.5× larger).

**Figure 2:**
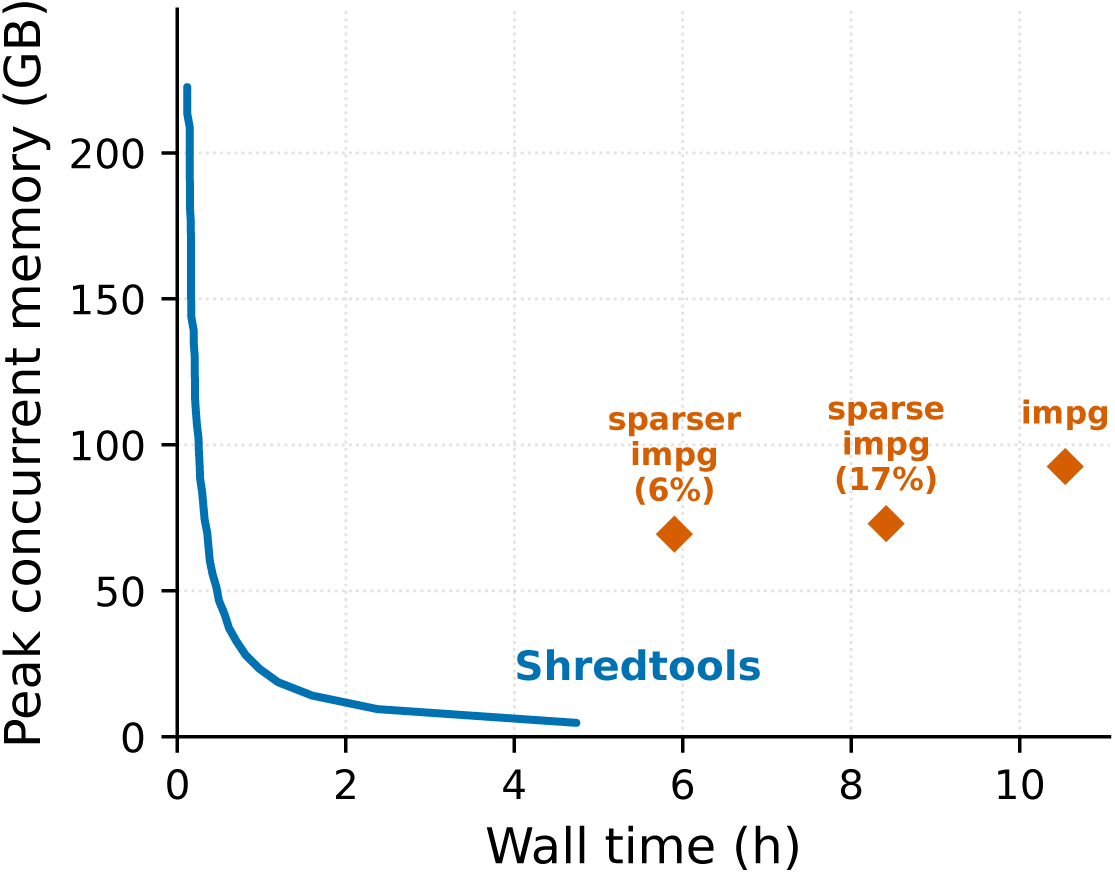
Comparison of time and memory used to construct a Shredtools multi-MUM index using Mumemto and an impg index using wfmash. wfmash was run using 48 threads. Mumemto was run with varying thread numbers to compare the time-memory tradeoff front.

We queried 1,000 random regions drawn from across the 92 chr19 genomes with both methods to compare speed and memory. Shredtools took an average of 0.484s and 79 MB peak memory footprint per query, compared to 4.01s and 1.66 GB for an impg query, an 8× speed increase. However, using the sparse set of pairwise alignments (17%, the default `auto` setting), impg queries improve to 0.21s and 12 MB of memory.

### Extract and enhance highlight local structural variation

We used Shredtools to compute the multi-MUM synteny of the *A. thaliana* pangenome (N=69) [23], as illustrated in the right side of Figure 3. To further demonstrate `extract`, we queried the pericentromeric region on the q-arm of chromosome 1, which contains multiple rearrangements and inversions [23, 24]. We queried the region `chr1:20425000-25500000` from the Db-1 genome and extracted coordinates for the syntenic regions in the other 68 genomes in less than one second. The extracted region had MUM bounds within 80 bp of either side, allowing the region to be extracted with minimal superfluous flanking sequence. We then augmented this view by running Shredtools’ `enhance` query on a subset of the genomes. This query iteratively re-runs Mumemto on gaps between collinear multi-MUMs; by doing so, we found additional MUMs that are also present in all genomes and which are unique, but their “uniqueness” is only with respect to the sequence in the gap between the globally-unique multi-MUMs. Computing these added a finer resolution to the region, especially near the centromere. This can be seen in Figure 3 left inset.

**Figure 3:**
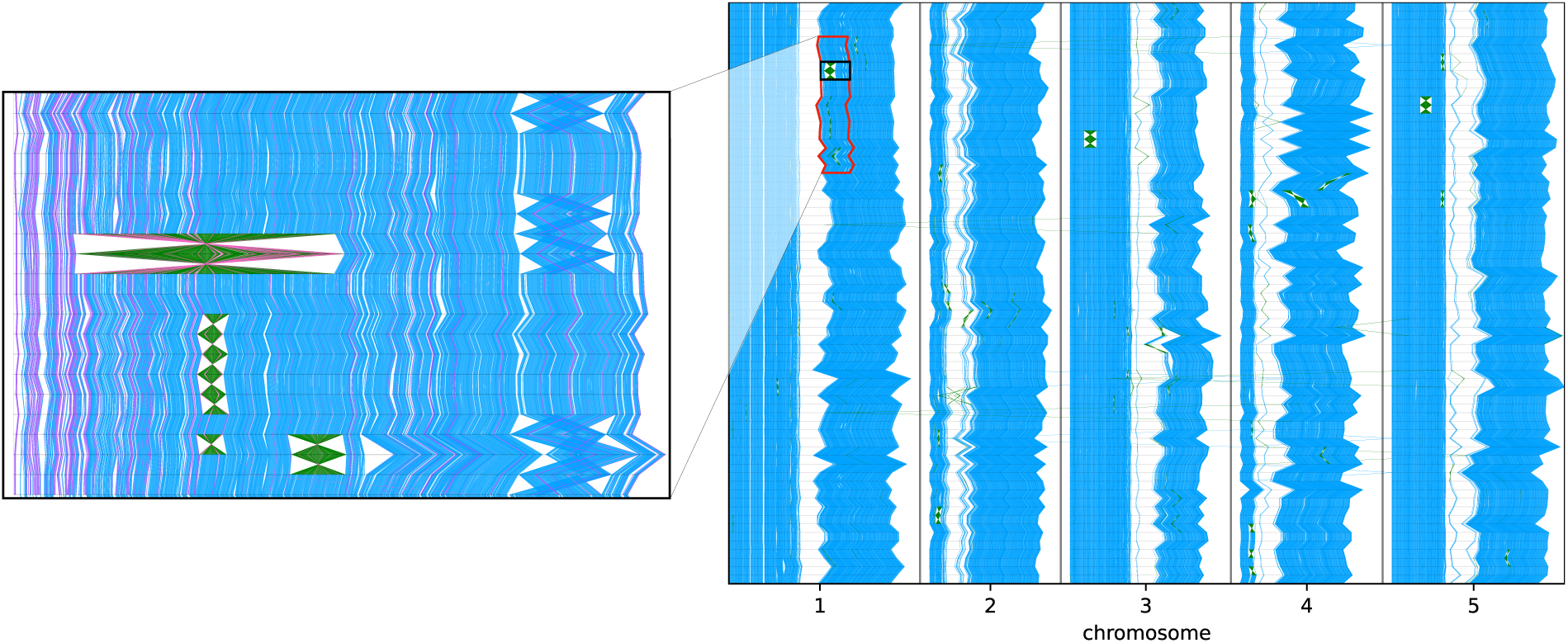
Multi-MUM synteny for 69 T2T *A. thaliana* genomes. An extract from the peri-centromeric region of chromosome 1 which exhibits multiple structural variants in a subset of genomes is shown on the left as a zoom panel. Local multi-MUMs from `enhance` are shown in purple (reverse strand matches in pink) to differentiate from strict multi-MUMs in blue (and green).

### Shredtools extracts genes from entire pangenomes

We applied `extract` to the human pangenome to assess how readily annotated gene loci can be retrieved across all genomes. We queried the 55,997 autosomal gene loci from the NCBI RefSeq annotation lifted to CHM13 v2.0 [13] using Liftoff [25]. Figure 4A shows a scatterplot of the `extract` bounds achieved, i.e. the distance to the nearest flanking multi-MUM on each side of a query, for all 55,997 gene regions. Of these, 34,864 (62.3%) had perfect bounds (Figure 4B), i.e. 0 on both sides (thus, at coordinate 0, 0 on the plot), meaning the query interval was inside and/or immediately flanked by multi-MUMs on both sides. An additional 4,404 genes (7.9%) had bounds within 100 bp on both sides. For medically relevant genes from the Online Mendelian Inheritance in Man (OMIM) collection [26], 67% were within 100 bp of a multi-MUM (Figure 4D), rising to 87% when restricting to the smaller HPRCr1 collection. This reflects greater multi-MUM density at smaller pangenome sizes, an important factor in deciding which pangenome to use.

**Figure 4:**
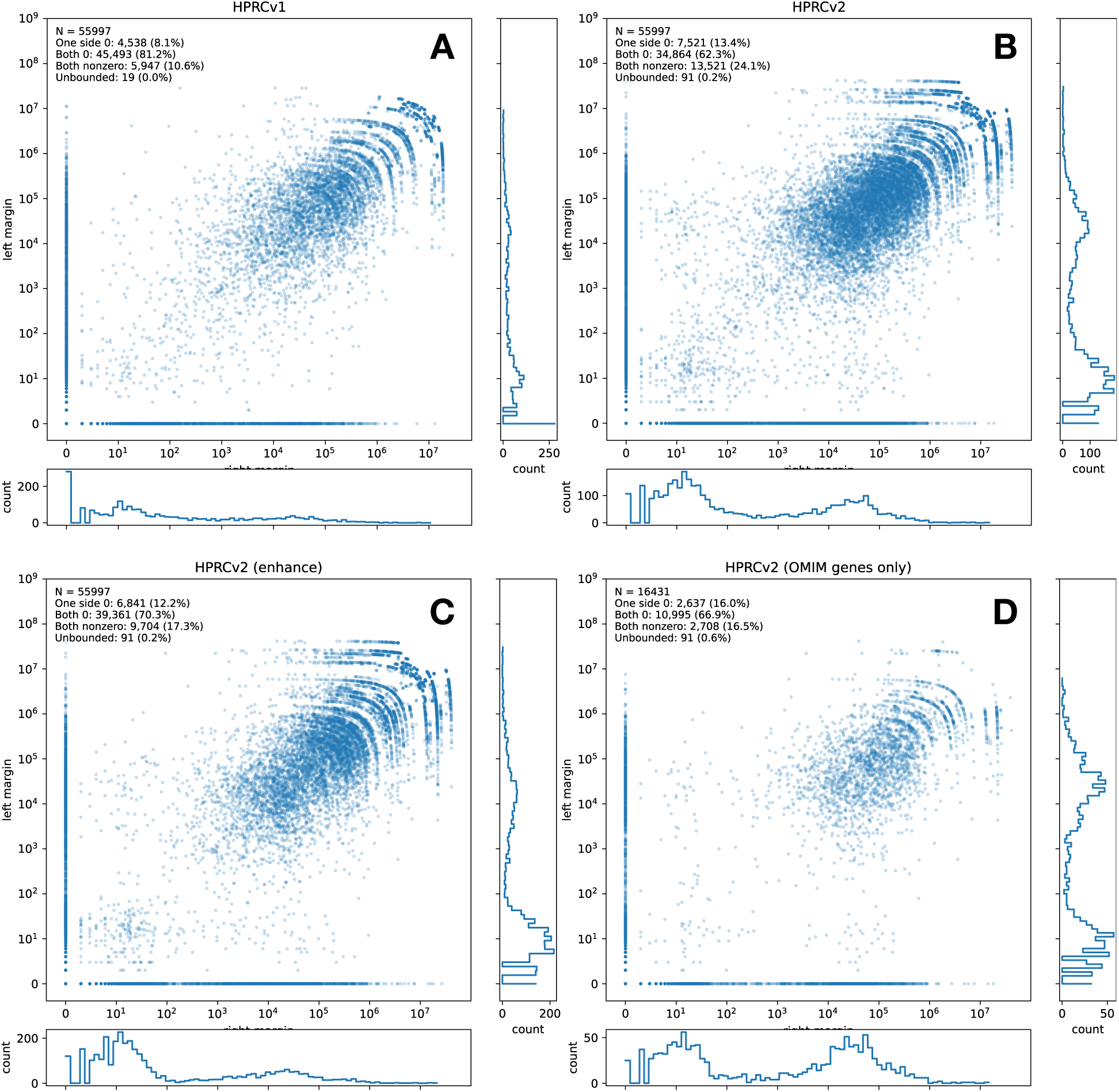
Comparison of `extract` bounds, the distance on each side of a query region to the nearest flanking multi-MUMs, for different Shredtools indexes. Histograms on the x and y-axis show the distribution of points with only one-sided zero bounds (shown as points in a line along the axis). Bounds tend to improve with higher multi-MUM cover-age, and `enhance` supplements the HPRCr2 set of multi-MUMs to improve resolution of `extract` for many regions.

#### Enhancing improves multi-MUM coverage

The `extract` bounds on a query interval are a function of the multi-MUM coverage in the region of interest. In areas where multi-MUMs have denser coverage, it is generally easy to understand and extract pangenomic regions of interest. The `enhance` function serves to increase multi-MUM coverage by finding *local* multi-MUMs, matches that are not globally unique, but are unique with respect to a gap between an adjacent pair of global multi-MUMs. Recursively searching for local multi-MUMs in such gaps was implemented in previous multi-MUM methods for similar reasons [27]. We applied the `enhance` strategy to the HPRCr2 dataset, finding local multi-MUMs to supplement the global coordinate system. For comparison, we computed the coverage of multi-MUMs across CHM13, i.e. the number of bases in the reference aligned to the pangenome as part of a multi-MUM. Across HPRCr1 genomes, 76% of the pangenome is covered by global multi-MUMs. This decreases as we add genomes, dropping to 63% for HPRCr2. After `enhance`, however, we gain an additional 7.6 percentage points in coverage.

As predicted, increasing coverage with local multi-MUMs also improves resolution of `extract` queries. When including local multi-MUMs for HPRCr2, we found that 4,497 gene regions that previously had imperfect bounds gained perfect bounds after `enhance`, a 13% increase from the strict multi-MUM pangenome for HPRCr2(Figure 4). Of those genes, 23 were considered challenging, medically-relevant genes [28]. One gene, MRC1, has been shown to be falsely duplicated in previous human genome assembly versions. Thus, this may be a case where the new MUMs introduced by `enhance` should have been globally unique as well, but became non-unique due to a misassembly in at least one sequence.

#### Enhance further refines multi-MUM bounds and improves interpretation

The Shredtools `enhance` feature also includes a second method, which improves accuracy of extractions even where multi-MUM coverage is sparse. In many cases, sparse multi-MUM coverage is due to the presence of structural variants or misassemblies in just a few (even just one) of the genomes in the pangenome. In such cases, we can improve our chance of extracting a relevant and tightly-bounded region by first discarding genomes that disagree with the global coordinates. Specifically, when attempting to extract a region, `enhance` will first enrich the region with local multi-MUMs, then it will also identify any *outlier* genomes in the local region, whose presence drives the local sparsity of multi-MUMs. Outliers are detected by examining the distribution of gap lengths between the flanking markers, since we observed that sparsity tends to be driven by genomes exhibiting aberrant inter-MUM gap lengths. We call this feature `zoom`, en-abling a user to zoom into local regions while being aware of outlier assemblies that may not conform to the local multi-MUM coordinate system.

As an example, we used Shredtools to extract the immunoglobulin kappa chain (IGK) V gene segment IGKV6-21 across 15 T2T reference genomes. Using annotations in CHM13 [29], we queried the region `chr2:89168754-89169041` and found very loose bounds: 328,599 bp and 74,147 bp to the left and right of the query region respectively. Running `enhance`, we found that two diploid genomes had particularly small extracted regions: HG002 and I002C. Both were derived from lymphoblastoid cell lines (LCLs), and thus contained VDJ recombination in the IGK locus which deletes many V gene segments. We used `enhance` to omit those two genomes and re-extracted the query region from the remaining genomes. This allowed us to obtain exact bounds (0 bp to the left and right) and thus perfectly extract the IGKV6-21 region from the remaining 13 genomes.

Iterative extract is also effective on regions that are duplicated, either through segmental duplication or misassembly. In these cases, explicitly omitting the outlier genome is often not required, as the extracted region tends to contain a single copy in each genome. Thus `enhance` and re-extracting can achieve better bounds within the local, single-copy context. One example of this scenario was the TERT gene on chr5, which encodes a component of the telomerase enzyme. Extracting this gene from HPRCr2 genomes yielded poor bounds, 25–35 Kbp on either side to the nearest marker; however incorporating local multi-MUMs resulted in perfect bounds. Computing partial multi-MUMs across the extracted local regions to find outliers, we found a single genome (paternal haplotype of HG04228) that was obfuscating the multi-MUMs in the region. Manually inspecting it, we found an alternate contig that duplicated the first 1.7 Mbp region of chr5. Despite this, by running `enhance` , we were able to overcome the misassembly and use the global coordinate system to extract the TERT gene precisely across all 476 genomes.

### A human pangenome web interface

The `bumbl.bi` index enables efficient queries in arbitrary regions of interest, framed in terms of the linear coordinates of any genome in the pangenome. Because the index stores byte offsets rather than coordinates, a remote client can answer a query by issuing a small number of HTTP Range requests directly against the `bumbl` file: one or two requests to read the relevant index windows, followed by three requests to fetch the corresponding multi-MUM rows from the LENGTHS, STARTS, and STRANDS sections of the `bumbl` file. For a typical flanking query over HPRCr2 ( 50 multi-MUM rows), the STARTS section read transfers about 190 KB. The total data transferred per query is therefore on the order of a few hundred kilobytes, regardless of the size of the underlying pangenome index.

We implemented this in a Pyodide web application that answers `extract` queries across the HPRCr1 or HPRCr2 pangenomes directly from a web browser (Figure 5). The multi-MUMs and `bumbl.bi` index are hosted in a static AWS S3 bucket; all query computation runs in the browser via WebAssembly, with no server-side compute required. This serverless architecture means the tool scales to arbitrary concurrent users with no operational infrastructure beyond storage. The user requests a region and receives a BED-file response listing the syntenic regions across the pangenome, together with the bounding distances to the nearest flanking multi-MUMs. A typical query takes 1–3 seconds depending on network latency. The user can optionally plot the synteny using a Pyodide WebAssembly version of the Mumemto multi-MUM plotting module, and download the extracted sequences from each HPRC assembly to a FASTA output.

**Figure 5:**
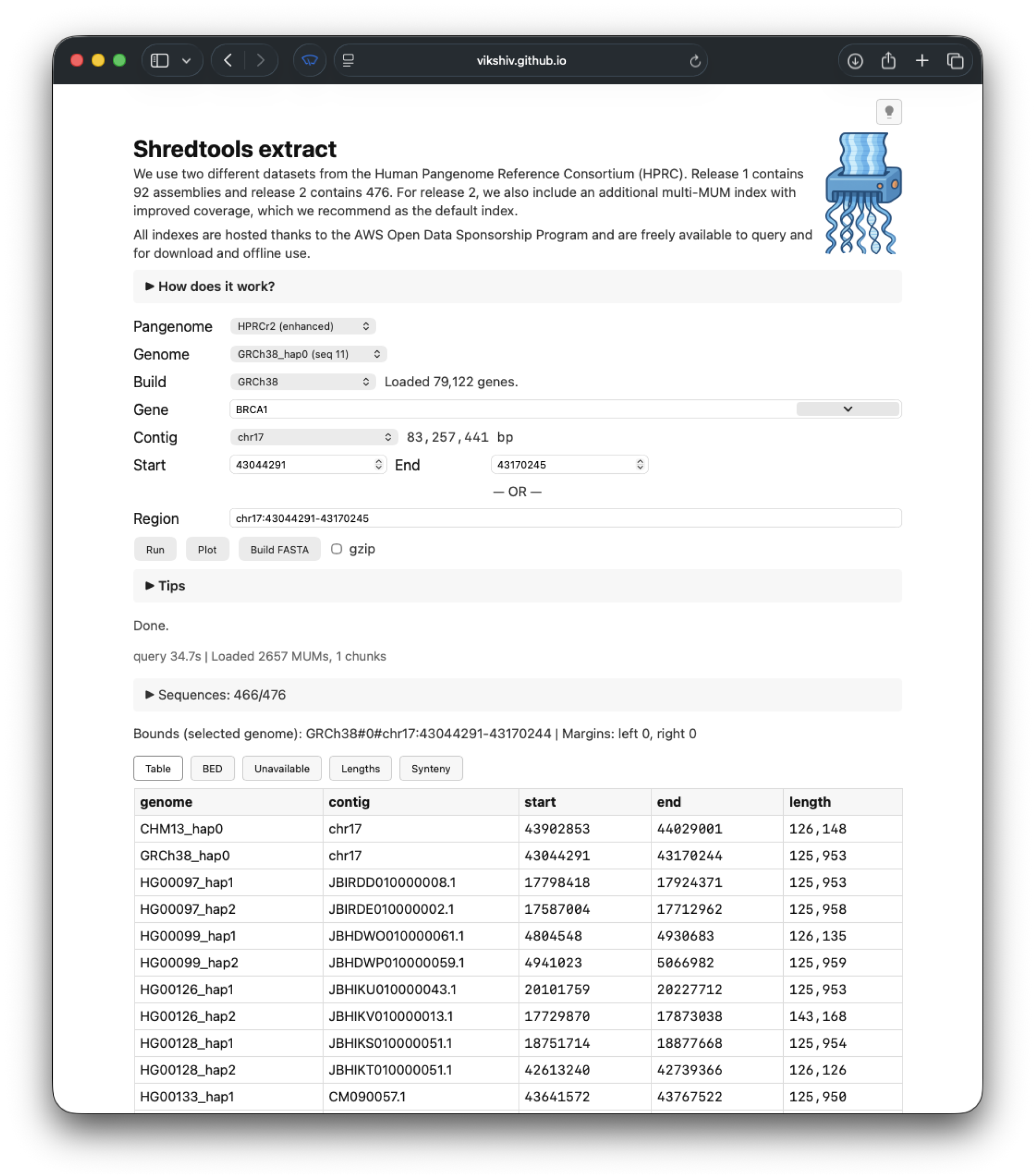
The Shredtools `extract` web interface. The user selects a human pangenome (HPRCr1 vs HPRCr2), then enters an interval with respect to any genome in the collection. Optionally, the user can enter a gene name for a gene present in the annotation for either CHM13 or GRCh38. Pangenome-wide syntenic regions are computed and displayed. The query computation happens via HTTP Range requests made to an index held in a public AWS bucket.

## Discussion

Shredtools demonstrates that a multi-MUM coordinate system—built without a full MSA or genome-scale pairwise alignment—is sufficient for sophisticated pangenome interval queries at scale. Using Shredtools, a user can extract syntenic sequences from a 476-genome pangenome in under a second, resolve regions with poor global coverage by zooming in on collinear subsets, or iteratively refine the coordinate system toward a fuller alignment, all using an index that remains up to date as new genomes are released.

We note that shredding can also be used to divide a pangenome into smaller, independent subunits for easier downstream computation. Recent work [30] developed a method to improve the efficiency of prefix-free parsing-based BWT construction by splitting the sequence into small, dissimilar regions and merging the resulting BWTs. Shredtools can supply the necessary non-overlapping regions—chosen to maximize intra-shred similarity and minimize inter-shred similarity—as input to this procedure.

A limitation of `extract` is its dependence on multi-MUM coverage: in regions where bases tend not to be covered by multi-MUMs, possibly due to one or more sequences carrying structural variants or misassemblies, the flanking markers may be distant and yield wide, imprecise bounds. Shredtools currently addresses this through zooming: iteratively narrowing the set of genomes to those for which a local coordinate system exists, then re-extracting on the reduced collection. The IGK locus case above illustrates this. That said, we note that zooming can be made still more systematic by leveraging Mumemto’s tree-based merging framework [31]. In this framework, intermediate multi-MUM computations are stored at each internal node of a partition hierarchy over the genomes; a query can then automatically select the node that includes the most genomes while also ensuring the target region is well covered by MUMs, without manual intervention.

We include a comparison with the impg tool, which performs similar queries to `extract` and `shred` to partition the pangenome into local regions for downstream analysis. We provide a benchmark of constructing an impg index over wfmash-computed pairwise alignments as an example. However, we note that impg is a general framework that can operate over different notions of pangenome homology. Indeed, a potential future direction would be to integrate the multi-MUM coordinate system into the impg framework.

The web interface for `extract` is powered by the `bumbl.bi` index, which enables efficient retrieval of data from a multi-MUM index either locally or over a network. The `bumbl.bi` format is analogous to BAM and BigBED indexes in this respect, suggesting natural integration with genome browsers such as IGV and JBrowse as a future direction; such integration would let users view multi-MUM synteny and run Shredtools queries interactively, alongside existing genomic annotations, in the same way that BAM indexes and BigBED files enabled interactive visualization of large genomic datasets.

Pangenome collections will continue to grow, and the MSA bottleneck that limits graph-based coordinate systems is unlikely to recede at the same pace. Multi-MUM-based coordinates offer a scalable alternative: they are easily computed and updated, they generalize fixed-length markers, and they support useful queries. Shredtools translates these properties into practical tools, and its integration with genome browsers, further application to nonhuman pangenomes, and tighter coupling with the Mumemto partition hierarchy are natural next steps as the field moves toward routine pangenome-scale analyses.

## Methods

### Mumemto

Shredtools queries rely on pre-computed maximal unique matches (multi-MUMs) computed using Mumemto [11]. A multi-MUM is an exact sequence match that appears in each genome in a collection exactly once (unique) and is non-extendable on either side (maximal). As of Mumemto v1.3, these matches can be computed in a dynamic, mergeable manner [31] using a partition-merge framework, making Mumemto applicable to pangenomes of practically any size. The resulting multi-MUM collection represents a dynamic index of conserved regions that can be updated as new genomes are released. Shredtools in particular uses collinear multi-MUMs, matches that are order-preserving neighbors across all genomes in a pangenome. These matches are more likely to represent conserved columns of the underlying MSA, and they partition the pangenome into distinct collinear units (shreds), each bounded by the same pair of multi-MUMs across all genomes.

The same Mumemto auxiliary structure also supports the `shred` comparison in Results. It stores, for every unique match across the collection, the longest prefix of that match that remains unique. Unique *k*-mers of any length *k* appear as positions where the auxiliary array value is nonzero and less than *k*, and the unique match at that position has length greater than *k*, enabling recovery of unique *k*-mer positions without a separate index [31].

**`bumbl.bi` index** Shredtools functions require efficient access to the collection of multi-MUMs that form the pangenome coordinate system. To facilitate this, we developed the `bumbl.bi` index. The `bumbl` format is Mumemto’s native binary format for multi-MUM collections; the `bumbl.bi` index adds a positional offset structure on top of it. The index has two formats: a single-index and a multi-index, enabling queries with respect to a single reference genome or any genome in the collection, respectively. Both formats support rapid retrieval of multi-MUMs in a local region without loading the full `bumbl` file into memory. This property is particularly important for network access, such as serving multi-MUM data from a cloudhosted `bumbl` file via HTTP Range requests.

### Single index

The single-index stores byte offsets into the `bumbl` file for non-overlapping windows across a chosen reference genome, analogously to a BAM index. It is fixed at indexing time and divided into 1 Mbp windows by default. For each window, the index records the position of the first multi-MUM that falls within it, in the sorted order of the `bumbl` file. In addition, each window stores two summary values: the minimum multi-MUM start position and the maximum multi-MUM end position within that window. These per-window spans allow the algorithm to determine whether a window can bound a query boundary without loading any multi-MUMs from the `bumbl` file. To answer a query, the algorithm identifies the nearest windows that can provide a left and right flank for the query interval (searching outward from each edge), retrieves the corresponding byte offsets, and fetches only those flanking multi-MUMs. Crucially, `extract` only ever needs the immediately flanking multi-MUMs on each side of the query, not all multi-MUMs in the interval. Thus, a typical query loads data from at most two windows regardless of how large the query region is.

### Multi index

The multiindex extends the format to support queries with respect to any genome. The `bumbl` file is first sorted by one member’s coordinates (typically the first sequence). Because Shredtools filters to collinear multi-MUMs, which are order-preserving across genomes by definition, their rank order in the sorted `bumbl` file is approximately preserved in every other genome’s coordinate system as well. Consequently, a genomic window in any genome maps to a small number of contiguous row intervals in the file. This compactness is a direct consequence of the collinearity filter: without it, a window in a non-reference genome could scatter across many disjoint row intervals. Each genome is divided into windows, and for each window the index stores the half-open row intervals together with the corresponding byte offsets and per-window span values. In practice, for HPRCr2 (N=476), nearly all windows correspond to a single contiguous interval, with at most three disjoint intervals, so the index remains compact despite covering all 476 genomes.

### extract

The `extract` function retrieves sequences homologous to a query region using collinear multi-MUMs, without requiring a sequence alignment. For a given query, the nearest collinear multi-MUMs flanking the region are identified and used to extract the enclosed sequences from the remaining genomes. The bounds depend on multi-MUM coverage in the region: how close the nearest flanking markers are to the query boundaries. These distances are reported to the user as guideline bounds.

### Iterative refinement

When multi-MUM coverage is low, the nearest flanking markers may be distant, producing loose bounds and an unclear picture of the pangenome. In such cases, an iterative process can tighten these bounds. After the initial extraction, multi-MUMs are re-computed over the extracted sequences from each genome. The resulting matches may not be globally unique, but they define a local coordinate system for the extracted region, enabling reextraction with tighter bounds. This process is repeated until the bounds stop improving or exact bounds are achieved.

### enhance

The `enhance` function improves multi-MUM coverage by finding local multi-MUMs in gaps between existing global multi-MUMs, thereby increasing the resolution of the pangenome coordinate system. Local multi-MUMs are matches that are locally unique within each genome’s gap region, but not necessarily globally unique. A similar recursive strategy for finding local MUMs was used in Parsnp [27] to improve core genome alignment coverage. These local matches still serve as coordinate markers, but may not be strict multi-MUMs due to segmental duplications, inverted repeats, or misassemblies. The `enhance` procedure enumerates all gaps between consecutive collinear multi-MUM pairs within each collinear block, then processes those where at least one genome has a gap of at least a user-supplied minimum length. For each processed gap, a genome is excluded if its gap length falls below median − 5 × MAD across genomes for that gap (the factor of 5 is configurable); this removes those with anomalously small gaps caused by private deletions or collapsed misassemblies. Mumemto is then run on the gap sequences from the rest to find local multi-MUMs, which are filtered to those that form collinear blocks within the extracted region and merged back into the global multi-MUM collection.

## Code availability

Shredtools is available open-source at https://github.com/vikshiv/shredtools and released under a GPL-3.0 license. Multi-MUMs were computed for each pangenome using Mumemto v1.4.0. The Shredtools browser tool is available publically at https:// vikshiv.github.io/shredtools/, and uses `bumbl.bi` indexes available publically at the “Index Zone” S3 bucket (s3://genome-idx). We used HPRCr2 assemblies available from https://human-pangenomics.s3.us-west-2.amazonaws.com.

## Authors’ contributions

V.S.S. and B.L. conceived the project. V.S.S. wrote the software and conducted the experiments. V.S.S. and B.L. wrote the manuscript. All authors revised and approved the final manuscript.

## Acknowledgments

We thank the AWS Open Data Sponsorship Program for sponsoring the “Index Zone” S3 bucket (`s3://genomeidx`), where the HPRC `bumbl.bi` indexes are hosted. We would like to acknowledge the National Human Genome Research Institute (NHGRI) for funding the following grants supporting the creation of the human pangenome reference: U41HG010972, U01HG010971, U01HG013760, U01HG013755, U01HG013748, U01HG013744, R01HG011274, and the Human Pangenome Reference Consortium (BioProject ID: PR-JNA730823).

## Collaborators

*Human Pangenome Reference Consortium Version 2 Authors:* Derek Albracht^1^, Ivan A. Alexandrov^2^, Jamie Allen^3^, Alawi A. Alsheikh-Ali^4^, Nicolas Altemose^5^, Casey Andrews^6^, Dmitry Antipov^7^, Lucinda Antonacci-Fulton^1^, Mobin Asri^8^, Marcelo Ayllon^9^, Jennifer R. Balacco^10^, Floris P. Barthel^11^, Edward A. Belter Jr^1^, Halle D. Bender^8^, Andrew P. Blair^8^, Davide Bolognini^12^, Katherine E. Bonini^13^, Christina Boucher^14^, Guillaume Bourque^15,16,17^, Silvia Buonaiuto^18^, Shuo Cao^18^, Andrew Carroll^19^, Ann M. Mc Cartney^8^, Monika Cechova^8^, Mark J.P. Chaisson^20^, Pi-Chuan Chang^19^, Xian Chang^8^, Jitender Cheema^3^, Haoyu Cheng^21^, Claudio Ciofi^22^, Hiram Clawson^8^, Sarah Cody^1^, Vincenza Colonna^18^, Holland C. Conwell^23^, Robert Cook-Deegan^24^, Mark Diekhans^8^, Maria Angela Diroma^22^, Daniel Doerr^25,26,27^, Zheng Dong^6^, Danilo Dubocanin^5^, Richard Durbin^28,29^, Jana Ebler^25,30^, Evan E. Eichler^9,31^, Jordan M. Eizenga^8^, Parsa Eskandar^8^, Eddie Ferro^14^, Anna-Sophie Fiston-Lavier^32,33^, Sarah M. Ford^23^, Willard W. Ford^34^, Giulio Formenti^10^, Adam Frankish^3^, Mallory A. Freeberg^3^, Qichen Fu^6^, Stephanie M. Fullerton^35^, Robert S. Fulton^1^, Shenghan Gao^36^, Yan Gao^37^, Gage H. Garcia^9^, Obed A. Garcia^38^, Joshua M.V. Gardner^8^, Shilpa Garg^39^, Erik Garrison^18^, Nanibaa’ A. Garrison^40,41,42^, John E. Garza^1^, Margarita Geleta^43^, Mohammadmersad Ghorbani^44^, Tina A. Graves-Lindsay^1^, Richard E. Green^23^, Carol W. Greider^45^, Cristian Groza^46^, Bida Gu^20^, Andrea Guarracino^11,18^, Melissa Gymrek^47^, Maximilian Haeussler^8^, Leanne Haggerty^3^, Ira M. Hall^48,49^, Nancy F. Hansen^7^, Yue Hao^11^, Mohammad Amiruddin Hashmi^4^, David Haussler^8^, Prajna Hebbar^8^, Peter Heringer^25,26,27^, Glenn Hickey^8^, Todd L. Hillaker^8^, S. Nakib Hossain^3^, Neng Huang^37,50^, Sarah E. Hunt^3^, Toby Hunt^3^, Alexander G. Ioannidis^5,8^, Nafiseh Jafarzadeh^8^, Nivesh Jain^10^, Erich D. Jarvis^10,31^, Maryam Jehangir^11^, Juan Jiang^6^, Eimear E. Kenny^13^, Juhyun Kim^7^, Bonhwang Koo^10^, Sergey Koren^7^, Milinn Kremitzki^1,6^, Charles H. Langley^51^, Ben Langmead^52^, Heather A. Lawson^6^, Daofeng Li^6^, Heng Li^37,50^, Wen-Wei Liao^48,49^, Jiadong Lin^9^, Tianjie Liu^6^, Glennis A. Logsdon^36^, Ryan Lorig-Roach^8^, Jonathan LoTempio Jr^53^, Hailey Loucks^8^, Jane E. Loveland^3^, Jianguo Lu^54^, Shuangjia Lu^48,49^, Julian K. Lucas^8^, Walfred Ma^20^, Juan F. Macias-Velasco^1,6,55^, Kateryna D. Makova^56^, Maximillian G. Marin^37,50^, Christopher Markovic^1^, Tobias Marschall^25,30^, Franco L. Marsico^18^, Fergal J. Martin^3^, Mira Mastoras^8^, Capucine Mayoud^32^, Brandy McNulty^8^, Jack A. Medico^10^, Julian M. Menendez^8^, Karen H. Miga^8^, Anna Minkina^57^, Matthew W. Mitchell^58^, Saswat K. Mohanty^59^, Younes Mokrab^44,60,61^, Jean Monlong^62^, Shabir Moosa^44^, Avelina Moreno-Ochando^63,64^, Shinichi Morishita^65^, Jonathan M. Mudge^3^, Katherine M. Munson^9^, Njagi Mwaniki^66^, Nasna Nassir^4^, Chiara Natali^22^, Shloka Negi^8^, Lingbin Ni^9^, Adam M. Novak^8^, Faith Okamoto^8^, Keisuke K. Oshima^36^, Pilar N. Ossorio^67^, Chie Owa^65^, Sadye Paez^10^, Benedict Paten^8^, Clelia Peano^12,68^, Adam M. Phillippy^7^, Brandon D. Pickett^7^, Laura Pignata^18^, Nadia Pisanti^66^, David Porubsky^9,69^, Pjotr Prins^18^, Timofey Prodanov^25,30^, Anandi Radhakrishnan^8^, T. Rhyker Ranallo-Benavidez^11^, Brian J. Raney^8^, Mikko Rautiainen^70^, Alessandro Raveane^12^, Andreas Rechtsteiner^45^, Luyao Ren^9,31^, Arang Rhie^7^, Fedor Ryabov^71,72^, Samuel Sacco^23^, Farnaz Salehi^18^, Michael C. Schatz^52,73^, Laura B. Scheinfeldt^74^, Aarushi Sehgal^34^, William E. Seligmann^23^, Mahsa Shabani^75^, Kishwar Shafin^19^, Shadi Shahatit^32^, Ruhollah Shemirani^13^, Vikram S. Shivakumar^52^, Swati Sinha^3^, Jouni Sirén^8^, Linnéa Smeds^59^, Steven J. Solar^7^, Marco Sollitto^10,22^, Nicole Soranzo^12,28,76^, Andrew B. Stergachis^9,57^, Marie-Marthe Suner^3^, Yoshihiko Suzuki^65^, Arda Sö ylev^25,30^, Ahmad Abou Tayoun^77,78^, Jack A.S. Tierney^3^, Chad Tomlinson^1^, Francesca Floriana Tricomi^3^, Mohammed Uddin^4,79^, Matteo Tommaso Ungaro^23,80^, Rahul Varki^14^, Flavia Villani^18^, Ivo Violich^8^, Mitchell R. Vollger^57^, Brian P. Walenz^7^, Charles Wang^81^, Lisa E. Wang^13^, Ting Wang^1,6,55^, Aaron M. Wenger^82^, Conor V. Whelan^10^, Zilan Xin^6^, Zheng Xu^6^, Kai Ye^83^, DongAhn Yoo^9^, Wenjin Zhang^6^, Ying Zhou^37^, Xiaoyu Zhuo^6^, Giulia Zunino^12^

*Affiliations:*

1. McDonnell Genome Institute, Washington University School of Medicine, St. Louis, MO 63108, USA
2. Department of Human Molecular Genetics and Biochemistry, Faculty of Medical and Health Sciences, Tel Aviv University, Tel Aviv 69978, Israel
3. European Molecular Biology Laboratory, European Bioinformatics Institute (EMBL-EBI), Wellcome Genome Campus, Hinxton, Cambridge CB10 1SD, UK
4. Center for Applied and Translational Genomics (CATG), Mohammed Bin Rashid University of Medicine and Health Sciences, Dubai Health, Dubai, UAE
5. Department of Genetics, Stanford University, Palo Alto, CA 94304 USA
6. Department of Genetics, Washington University School of Medicine, St. Louis, MO 63110, USA
7. Genome Informatics Section, Center for Genomics and Data Science Research, National Human Genome Research Institute, National Institutes of Health, Bethesda, MD 20892, USA
8. UC Santa Cruz Genomics Institute, University of California, Santa Cruz, CA 95060, USA
9. Department of Genome Sciences, University of Washington School of Medicine, Seattle, WA 98195, USA
10. The Vertebrate Genome Laboratory, The Rockefeller University, New York, NY 10065, USA
11. Bioinnovation and Genome Sciences, The Translational Genomics Research Institute (TGen), Phoenix, AZ 85004, USA
12. Human Technopole, Milan, Italy
13. Institute for Genomic Health, Icahn School of Medicine at Mount Sinai, New York, NY 10029, USA
14. Department of Computer and Information Science and Engineering, University of Florida, Gainesville, FL 32611, USA
15. Canadian Center for Computational Genomics, McGill University, Montréal, QC H3A 0G1, Canada
16. Department of Human Genetics, McGill University, Montréal, QC H3A 0G1, Canada
17. Victor Phillip Dahdaleh Institute of Genomic Medicine, Montréal, QC H3A 0G1, Canada
18. Department of Genetics, Genomics and Informatics, University of Tennessee Health Science Center, Memphis, TN 38163, USA
19. Google LLC, Mountain View, CA 94043, USA
20. Quantitative and Computational Biology, University of Southern California, Los Angeles, CA 90089, USA
21. Department of Biomedical Informatics and Data Science, Yale School of Medicine, New Haven, CT 06510, USA
22. Department of Biology, University of Florence, Sesto Fiorentino, FI 50019, Italy
23. Department of Ecology and Evolutionary Biology, University of California, Santa Cruz, CA 95060, USA
24. Arizona State University, Consortium for Science, Policy & Outcomes, Washington, DC 20006, USA
25. Center for Digital Medicine, Heinrich Heine University Düsseldorf, Düsseldorf, NRW, DE
26. Department for Endocrinology and Diabetology at the Medical Faculty and University Hospital Düsseldorf, Heinrich Heine University Düsseldorf, Düsseldorf, NRW, DE
27. Paul-Langerhans-Group Computational Diabetology, German Diabetes Center (DDZ) and Leibniz Institute for Diabetes Research, Düsseldorf, NRW, DE
28. Wellcome Sanger Institute, Genome Campus, Hinxton, CB10 1RQ, UK
29. Department of Genetics, University of Cambridge, Cambridge, CB2 3EH, UK
30. Institute for Medical Biometry and Bioinformatics, Medical Faculty and University Hospital Düsseldorf, Heinrich Heine University, Düsseldorf, NRW, DE
31. Howard Hughes Medical Institute, Chevy Chase, MD 20815, USA
32. ISEM, Univ Montpellier, CNRS, IRD, Montpellier, FR
33. Institut Universitaire de France, Paris, FR
34. Department of Computer Science and Engineering, University of California San Diego, La Jolla, CA 92093, USA
35. Department of Bioethics & Humanities, University of Washington School of Medicine, Seattle, WA 98195, USA
36. Department of Genetics, Epigenetics Institute, Perelman School of Medicine, University of Pennsylvania, Philadelphia, PA 19104, USA
37. Department of Data Science, Dana-Farber Cancer Institute, Boston, MA 02215, USA
38. Department of Anthropology, University of Kansas, Lawrence, KS 66045, USA
39. School of Health Sciences, University of Manchester, Manchester M13 9PL, UK
40. Traditional, ancestral and unceded territory of the Gabrielino/Tongva peoples, Institute for Society & Genetics, University of California, Los Angeles, Los Angeles, CA 90095, USA
41. Traditional, ancestral and unceded territory of the Gabrielino/Tongva peoples, Institute for Precision Health, David Geffen School of Medicine, University of California, Los Angeles, Los Angeles, CA 90095, USA
42. Traditional, ancestral and unceded territory of the Gabrielino/Tongva peoples, Division of General Internal Medicine & Health Services Research, David Geffen School of Medicine, University of California, Los Angeles, Los Angeles, CA 90095, USA
43. Department of Electrical Engineering and Computer Science, University of California, Berkeley, Berkeley, CA 94720, USA
44. Medical and Population Genomics Lab, Sidra Medicine, Doha, Qatar
45. Department of Molecular Cell and Developmental Biology, University of California, Santa Cruz, CA, USA
46. Montreal Heart Institute, Montréal, QC, Canada
47. Department of Pediatrics, University of California San Diego, La Jolla, CA 92093, USA
48. Center for Genomic Health, Yale University School of Medicine, New Haven, CT 06510, USA
49. Department of Genetics, Yale University School of Medicine, New Haven, CT 06510, USA
50. Department of Biomedical Informatics, Harvard Medical School, Boston, MA 02115, USA
51. Department of Evolution and Ecology and the Center for Population Biology, University of California, One Shields, Davis, CA 95616, USA
52. Department of Computer Science, Johns Hopkins University, Baltimore, MD 21218, USA
53. Department of Pediatrics, Division of Genetics, School of Medicine, University of California, Irvine, CA 92697, USA
54. Sun Yat-sen University, Guangzhou, China
55. Edison Family Center for Genome Sciences & Systems Biology, Washington University School of Medicine, St. Louis, MO 63110, USA
56. Department of Biology and Center for Medical Genomics, Penn State University, University Park, PA 16802, USA
57. Division of Medical Genetics, Department of Medicine, University of Washington School of Medicine, Seattle, WA 98195, USA
58. The Jackson Laboratory for Genomic Medicine, Farmington, CT 06032, USA
59. Department of Biology, Penn State University, University Park, PA 16802, USA
60. Department of Biomedical Science, College of Health Sciences, Qatar University, Doha, Qatar
61. Department of Genetic Medicine, Weill Cornell Medicine-Qatar, Doha, Qatar
62. IRSD - Digestive Health Research Institute, University of Toulouse, INSERM, INRAE, ENVT, UPS, Toulouse, FR
63. MATCH biosystems, S.L., Elche, Spain
64. Universidad Miguel Hernández de Elche, Elche, Spain
65. Department of Computational Biology and Medical Sciences, The University of Tokyo, Kashiwa, Chiba 277-8561, Japan
66. Department of Computer Science, University of Pisa, Pisa, Italy
67. Law School, University of Wisconsin-Madison, Madison, WI 53706, USA
68. Institute of Genetics and Biomedical Research, UoS of Milan, National Research Council, Milan, Italy
69. Genome Biology Unit, European Molecular Biology Laboratory (EMBL), Heidelberg, DE
70. Institute for Molecular Medicine Finland, Helsinki Institute of Life Science, University of Helsinki, Helsinki, Finland
71. The Center for Bio- and Medical Technologies, Moscow, RUS
72. Centre for Biomedical Research and Technology, HSE University, Moscow, RUS
73. Department of Biology, Johns Hopkins University, Baltimore, MD 21218, USA
74. Coriell Institute for Medical Research, Camden, NJ 08103, USA
75. University of Amsterdam, Amsterdam, Netherlands
76. School of Clinical Medicine, University of Cambridge, Cambridge, CB2 0SP, UK
77. Center for Genomic Discovery, Mohammed Bin Rashid University, Dubai Health, UAE
78. Dubai Health Genomic Medicine Center, Dubai Health, UAE
79. GenomeArc Inc, Mississauga, ON, Canada
80. Department of Biology and Biotechnologies “Charles Darwin”, University of Rome “La Sapienza”, Rome 00185, IT
81. Center for Genomics, Loma Linda University School of Medicine, Loma Linda, CA 92350, USA
82. PacBio, Menlo Park, CA 94025, USA
83. The first affiliated hospital of Xi’an Jiaotong University, Xi’an Jiaotong University, Xi’an, Shaanxi, 710049, China

